# Sequencing of Sars-CoV-2 genome using different Nanopore chemistries

**DOI:** 10.1101/2021.01.02.425072

**Authors:** Oscar González-Recio, Mónica Gutiérrez-Rivas, Ramón Peiró-Pastor, Pilar Aguilera-Sepúlveda, Cristina Cano-Gómez, Miguel Ángel Jiménez-Clavero, Jovita Fernández-Pinero

## Abstract

Nanopore sequencing has emerged as a rapid and cost-efficient tool for diagnostic and epidemiological surveillance of SARS-CoV-2 during the COVID-19 pandemic. This study compared results from sequencing the SARS-CoV-2 genome using R9 vs R10 flow cells and Rapid Barcoding Kit (RBK) vs Ligation Sequencing Kit (LSK). The R9 chemistry provided a lower error rate (3.5%) than R10 chemistry (7%). The SARS-CoV-2 genome includes few homopolymeric regions. Longest homopolymers were composed of 7 (TTTTTTT) and 6 (AAAAAA) nucleotides. The R10 chemistry resulted in a lower rate of deletions in timine and adenine homopolymeric regions than R9, at expenses of a larger rate (~10%) of mismatches in these regions.

The LSK had a larger yield than RBK, and provided longer reads than RBK. It also resulted in a larger percentage of aligned reads (99% vs 93%) and also in a complete consensus genome.

The results from this study suggest that the LSK used on a R9 flow cell could maximize the yield and accuracy of the consensus sequence when used in epidemiological surveillance of SARS-CoV-2.

**Keypoints:**

- Sequencing SARS-CoV-2 genome is of great importance for the pandemic surveillance
- Nanopore offers a low cost and accurate method to sequence SARS-CoV-2 genome
- Ligation sequencing is preferred rather than the rapid kit using transposases

## INTRODUCTION

The human pathogen severe acute respiratory syndrome coronavirus 2 (SARS-CoV-2) emerged in Wuhan City (China) in early 2020 and causes the COVID-19 disease which is responsible for over one million deaths in less than one year since then. SARS-CoV-2 has spread over the world and has caused marked social, medical and economical adaptations to the world. Sequencing the genome of the SARS-CoV-2 has provided relevant information on the zoonotic origin of the virus (ref), its spreading dynamic, or its mutation rate. Genomic surveillance of SARS-CoV-2 is a key tool to know which lineages of the virus are circulating in the each country, how often new sources of virus are introduced from other geographical areas, or as an indicator of the success of control measures, and how the virus evolves in response to interventions. Sequencing also provide invaluable insights when linked with detailed epidemiology data for epidemiological investigation of the evolution of the pandemic. All these aspects play a key role in surveillance of the pandemic. Joint efforts have contributed to create a nomenclature system for the different linages of the coronavirus (Rambaut et al. 2020).

Recent documentation of reinfections has been provided demonstrating that different lineages of the SARS-CoV-2 can infect the same person (Tillett et al. 2020; To et al. 2020). Sequencing the genome of the coronavirus is necessary to confirm those reinfections and exclude medical relapses.

Fast and reliable sequencing of samples in hospitals is of main importance for this epidemiological surveillance. Oxford Nanopore Technologies (ONT) has developed several strategies for fast sequencing the SARS-CoV-2 genome, that may be essential for quick diagnosis and monitoring the community transmission of the coronavirus.

The objective of this study was to evaluate the accuracy of ONT sequencing of the SARS-CoV-2 genome using two chemistry and two sequencing strategies.

## MATERIALS AND METHODS

### Sample collection

A panel of clinical samples obtained during initial diagnostics in essential personnel from Madrid city hall services (police, firemen, emergency and health care workers, etc) was included in this study. Specifically, 10 nasopharyngeal swabs confirmed as SARS-CoV-2 positive by an in-house version of the recommended Egene real-time RT-PCR (Corman et al. 2020) (19.24≤Ct≤35) were selected. RNA was isolated anew from stored clinical samples using the IndiSpin®Pathogen kit (Indical Bioscience). Obtaining the cDNA and 400bp amplicons was conducted using protocol published by the ARTIC Network (Quick 2020), using primers from V2. The sample with the largest DNA concentration (15ng/μl) measured using the Qubit fluorometer (ThermoFisher Scientific, 150 Waltham, MA, USA) was selected for sequencing.

### DNA sequencing

The MinION device was used for ONT sequencing. Two ONT kits were used to prepare the DNA library: The Rapid Barcoding kit (SQK-RBK004) and the Ligation Sequencing Kit (SQK-LSK109).

The first library was sequenced using a R9.4 flow cell by loading 175 ng onto the SpotON flow cell, according to the manufacturer’s instruction. The other library was sequenced using the R10 flow cell using the same amount of DNA. Both flow cells (FC) ran until exhaustion. Most of the reads were obtained during the first two hours of the run. The flow cells were controlled and monitored using the MinKNOW software.

Reads were basecalled using Guppy version 4.0.11 (<u>community.nanoporetech.com</u>), and the high accuracy version of the flip-flop algorithm.

### Data analysis

The FASTQ files were aligned against the NCBI-nr protein database (Nov. 2017) using DIAMOND v0.9.22 (blastx option) setting the -F option to 15, to consider frame-shift errors in the sequences and the -range Culling and -top options set to 10 to scan the whole sequence for alignments with a 10% of the best local bit score (<u>megan.informatik.uni-tuebingen.de</u>, accessed on October 2018). The taxonomic binning of short reads from Illumina was performed using the daa2rma program from MEGAN Community Edition (CE) v6.11. with the option -a2t, to map the reads to the NCBI-taxonomy mapping file containing protein accessions (May 2017). Long reads from ONT were analyzed with specific parameters for long reads -lg (long reads) and -alg set to “longReads” (Huson et al. 2018). Relative taxonomic abundances were obtained for each samples and platform representing the number of reads assigned to each taxon. Relative abundances, alpha (Shannon and Simpson indexes) and beta diversity -using Bray-Curtis dissimilarity-were analyzed using the phyloseq (McMurdie and Holmes 2013), vegan (Oksanen et al. 2019), and microbiome R (Leo Lahti et al., 2012-2019) packages.

### Assembly

The SARS-CoV-2 virus from Wuhan strain Hu-1 genome (MN908947) was used as reference. The reads were aligned against SARS-CoV-2 genome using Minimap2 aligner (Li 2016), a general purpose alignment program to map DNA or long mRNA sequences against a reference database.

The total coverage of the genome for both sets of reads was calculated from the alignments using GenomeCoverageBed utility of the bedtools suite (Quinlan and Hall 2010), quantile-normalized and smoothed using a window width of 200bp.

### Variant calling

The variant calling genotyping from alignment files was performed using LoFreq (Wilm et al. 2012), VarScan (Koboldt et al. 2012) and Pilon (Walker et al. 2014). LoFreq is a fast and sensitive variant-caller for inferring SNVs and indels from NGS data. It makes full use of base-call qualities and other sources of errors inherent in sequencing. VarScan employs a robust heuristic/statistic approach to call variants that meet desired thresholds for read depth, base quality, variant allele frequency, and statistical significance. This program needs to preprocess the alignment file to generate a MPILEUP format file. Pilon identifies small variants with high accuracy as compared to state-of-the-art tools and is unique in its ability to accurately identify large sequence variants including duplications and resolve large insertions. Deviations from the reference were analyzed.

### De-novo assembly

The *de novo* assembly was performed using Canu (Koren et al. 2017). Canu is a fork of the Celera Assembler designed for high-noise single-molecule sequencing (such Oxford Nanopore MinION). In order to improve de genome assembly, we used Pilon to automatically improve draft assemblies. Pilon requires as input a FASTA file of the assembly along with the BAM files of reads aligned to the input FASTA file. Pilon uses read alignment analysis to identify inconsistencies between the input assembly and the evidence in the reads. The new assembled genomes were compared with reference genome using Gepard (Krumsiek *et al*., 2007) in order to thoroughly check the new assemblies.

## RESULTS

### Quality check

After the QC for ONT reads, a total of 16991 (N50=409) and 9658 (N50=1059) good quality reads (Table 1) were retained from the R9 and R10 FC, respectively. The R9 yielded 6.48 Mb and the R10 7.73Mb. ONT runs may yield much larger number of bases, however, in this case the amount of DNA was limiting. The GC content distribution were computed from both runs using LongQC (Fukasawa *et al*., 2020). The SARS-CoV2 reference genome has a GC% of 37.97. Reads from R9 averaged a GC content of 39.99% (s.d.=3.134), whereas GC content from R10 reads averaged 37.91% (s.d.=3.30). Fig. 1 represents the GC fraction in reads obtained from each FC. The entire reads are expected to show sharper distribution, because they have smaller deviation due to longer sequences. Red one is more robust to sequencing or sample differences, and this should be comparable to other data if the same target (biological replicates) is sequenced.

**Fig. 1.**
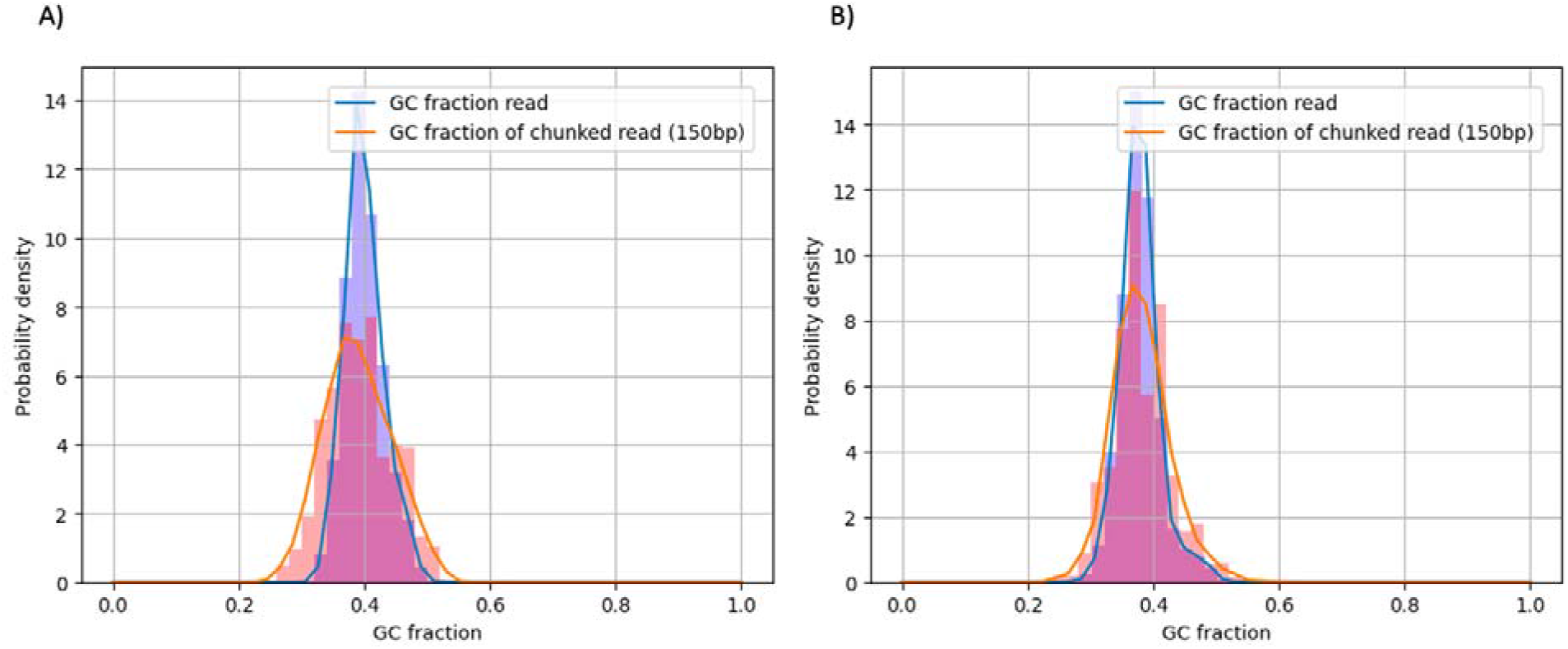
GC content distribution from ONT sequences from A) R9 set B) R10 flow-cells. The blue bars come from entire reads, and the red ones were computed from chunked (150bp) subsequences.

**Table 1.**
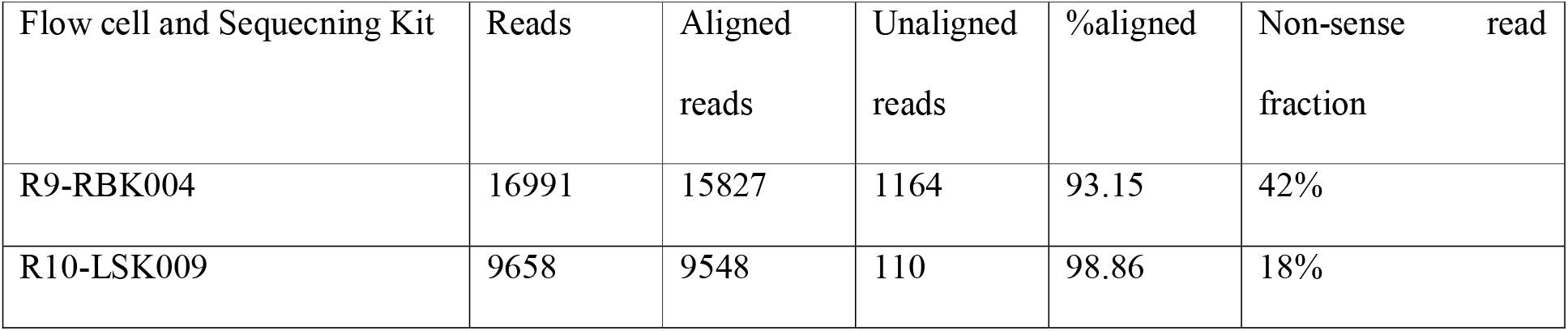
Minimap2 alignment summary results

| Flow cell and Sequecning Kit | Reads | Aligned<br>reads | Unaligned<br>reads | %aligned | Non-sense read<br>fraction |
| --- | --- | --- | --- | --- | --- |
| R9-RBK004 | 16991 | 15827 | 1164 | 93.15 | 42% |
| R10-LSK009 | 9658 | 9548 | 110 | 98.86 | 18% |

### Alignment

Brief statistics of the alignments are shown in Table 1. A larger proportion (98.86%) of reads from R10 were aligned to the reference genome. Almost 7% of reads from R9 were not aligned to the reference genome vs only 1.1% from R10 reads. The alignments show a good quality of reads for the process. However, a larger amount of non-sense reads (Fukasawa *et al*., 2020) were detected from R9 FC. Nonsense reads are defined as unique reads that cannot be mapped onto sequences of any other molecules in the same library. This concept is similar to unmappable reads; however, mappability depends on references. According to Fukasawa *et al*. (2020), non-sense read fraction should be less than 30%. If the fraction of non-sense reads is a way high, it might indicate that either sequencing had some issues or simply coverage is insufficient.

Fig. 2 shows the mismatches per aligned read against the SARS-CoV2 reference genome. The mismatches distributions showed a mode of 3.5% and 7% of read length for R9 and R10, respectively. The R9 FC showed a lower rate of mismatches than R10, although it might be led by a shorter length of the reads, which is a consequence of Rapid Barcoding kit (SQK-RBK004) used to prepare this library, as this kit requires a transposase fragmentation. However, the library loaded in the R10 FC was prepared using the Ligation Sequencing kit (SQK-LSK109) which does not fragment the DNA and is optimized for throughput. This might explain the slightly larger yield outcome from the library loaded in the R10 FC.

**Fig. 2.**
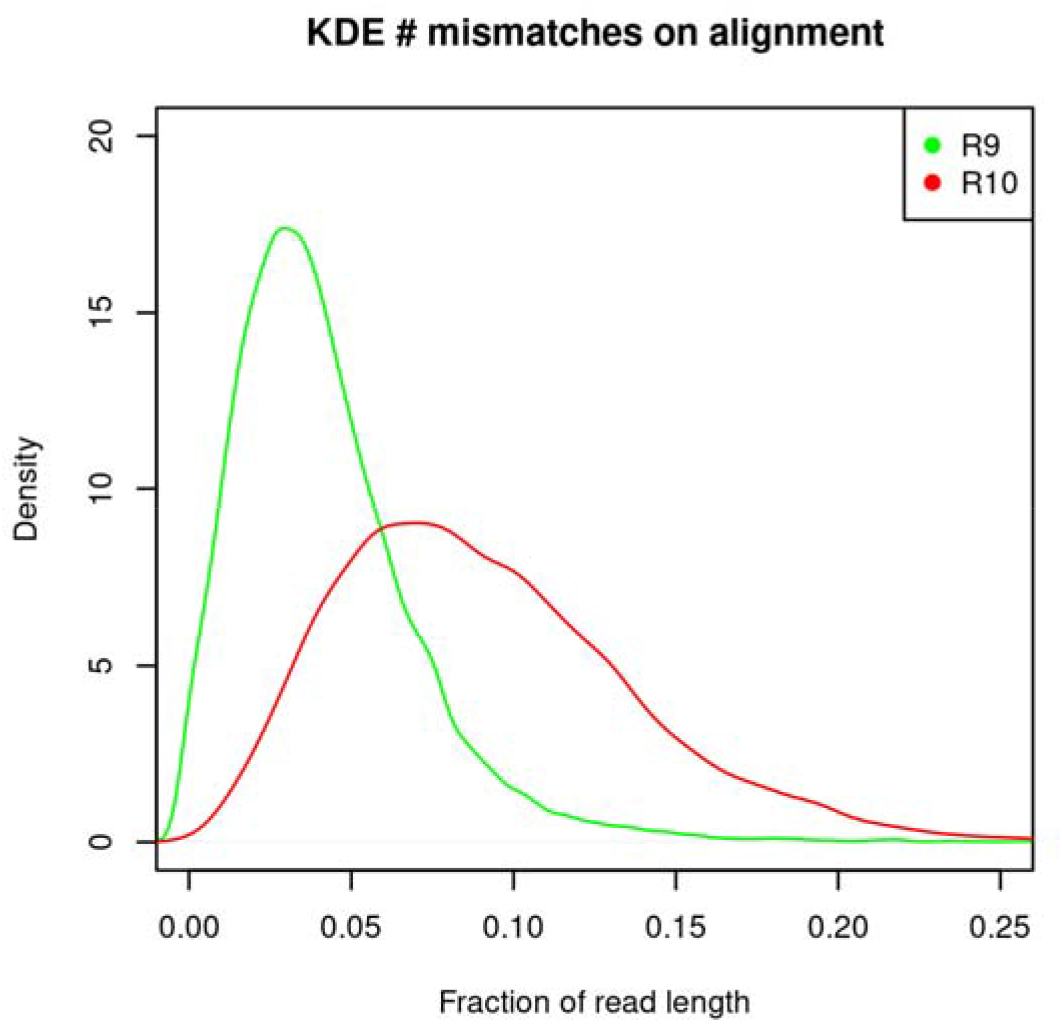
Fraction of mismatches per read against the SARS-CoV2 reference genome.

The read accuracy at the homopolymeric sites from each FC was evaluated. Homopolymeric sites in the SARS-CoV-2 reference genome were located with SeqKit (Shen *et al*., 2016). Longer homopolymeric regions in the SARS-Cov-2 reference genome were composed of timine (7 nucleotides) and adenine (6 nucleotides), whereas guanine and citosine homopolymers had a longest region of 3 nucleotides. Mismatches (Fig. 3A) and deletions (Fig. 3B) at homopolymer sites with length > 2 were considered. The R9 chemistry produced a lower number of mismatches at the homopolymeric sites, but was more prone than R10 to produce deletions, mainly in timine and adenine homopolymeric sites. The R10 chemistry produced a larger rate of mismatches, but a lower rate of deletions, although a rate >20% of deletions was observed at homopolymers <4 nucleotides. Both chemistries showed larger accuracy in adenine and timine homopolymeric sites than in citosine and guanine sites.

**Fig. 3.**
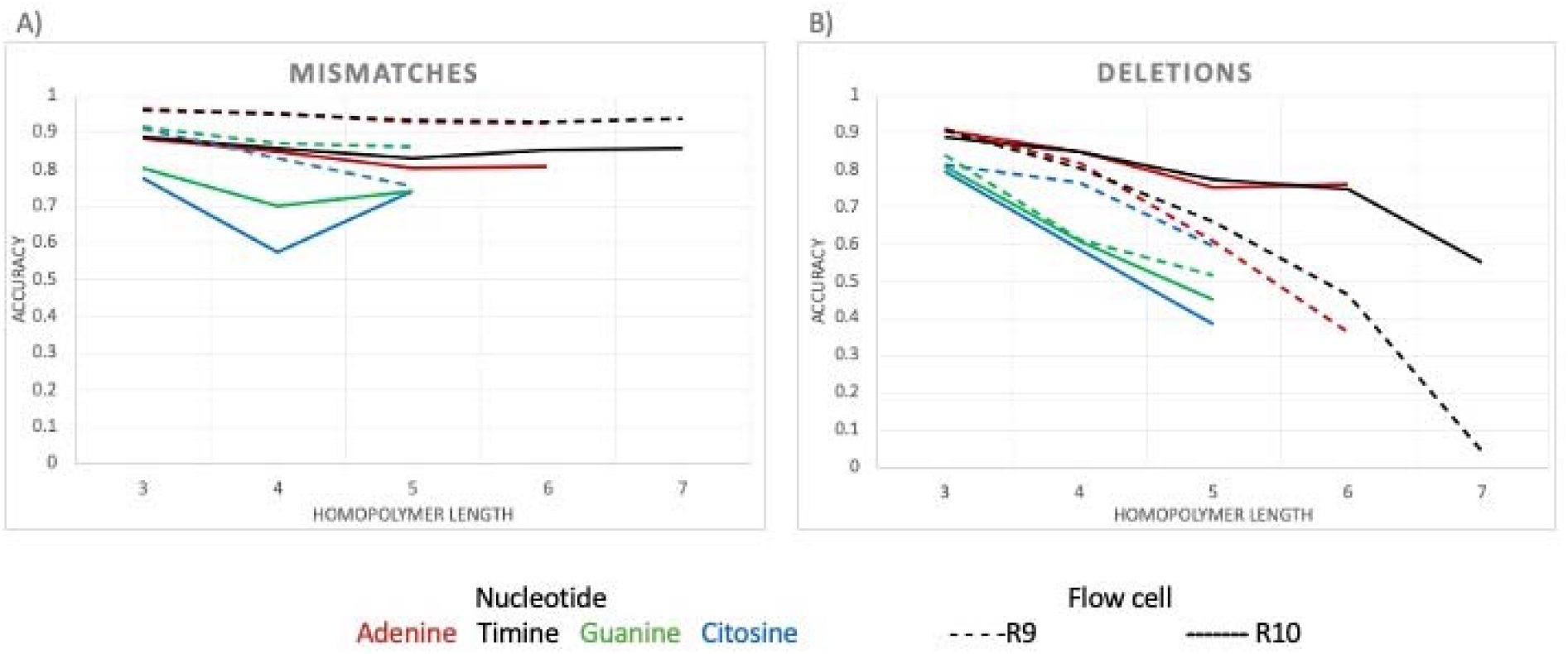
Read accuracy at homopolymeric sites in the SARS-Cov-2 genome for each FC type.

### Genome Coverage

Fig. 4 shows the log2 normalized smoothed coverage plus 1 (to avoid zeroes) plot, generated using GNUPLOT, to avoid zeroes. Most regions showed a coverage above 50x, and a coverage >200x was obtained for many regions, regardless the FC type. Both FC types had lower coverage in the same regions. It denotes that the primers used (ARTIC V2) produce a good coverage of the whole SARSCoV2 genome, although the 19k-20k region and the ending region showed a lower coverage than the rest of the genome.

**Fig. 4.**
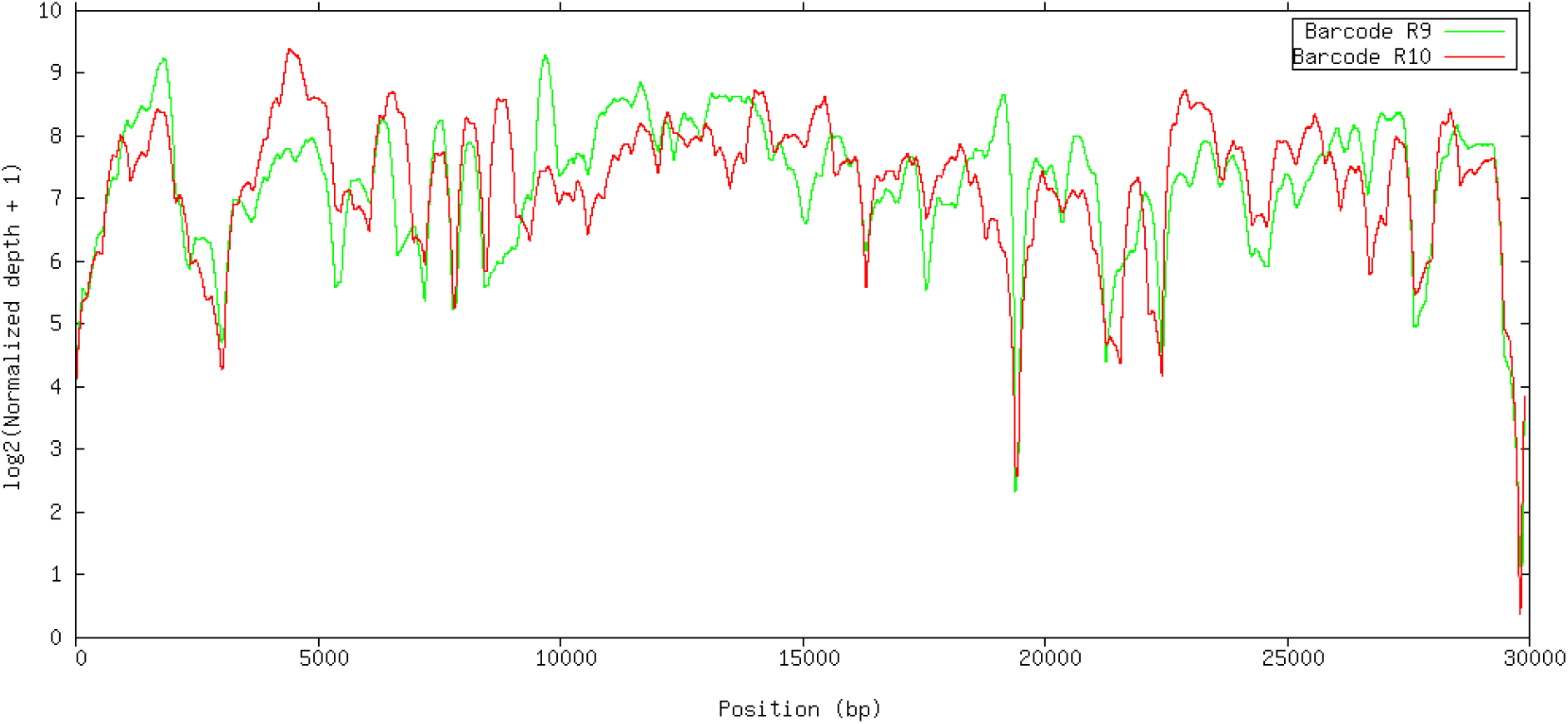
Smoothed normalized coverage of reads by position from each type of FC (R9 and R10) against the SARS-CoV-2 genome (smoothing window width = 200bp).

### Variant calling

Indels and SNP variants were identified using three softwares: LoFreq, Pilon and VarScan. There was a large variability for the number of detected variants between different programs. LoFreq detected a larger number of SNP variants from R9 (25 vs 9), Pilon reported similar number of SNPs for both FC (6 vs 7), whereas VarScan detected larger number of SNPs from R10 (35 vs 12). Common SNPs detected in both FC were consistent, with 6-7 SNPs detected in common in both FC in all the analyses. Fig. 5 and 6 show the Venn diagrams of common SNPs by FC and software, respectively. The common SNP variants detected were in frequency >0.5, as shown in Fig. 7. Requiring a large frequency (>0.60) of the detected variants from Nanopore long reads was consistent with the SNP variants detected from several software. Eight indels were reported in common from R9 and R10, however none of them were found with frequency > 0.5 (Fig. 8). Forty-seven other indels were detected only from R9 FC, and 31 indels only from R10. The combination of these strategies (i.e. large variant frequency and selecting common variants reported from different software) seems to be a reliable strategy for variant calling from error prone long reads.

**Fig. 5.**
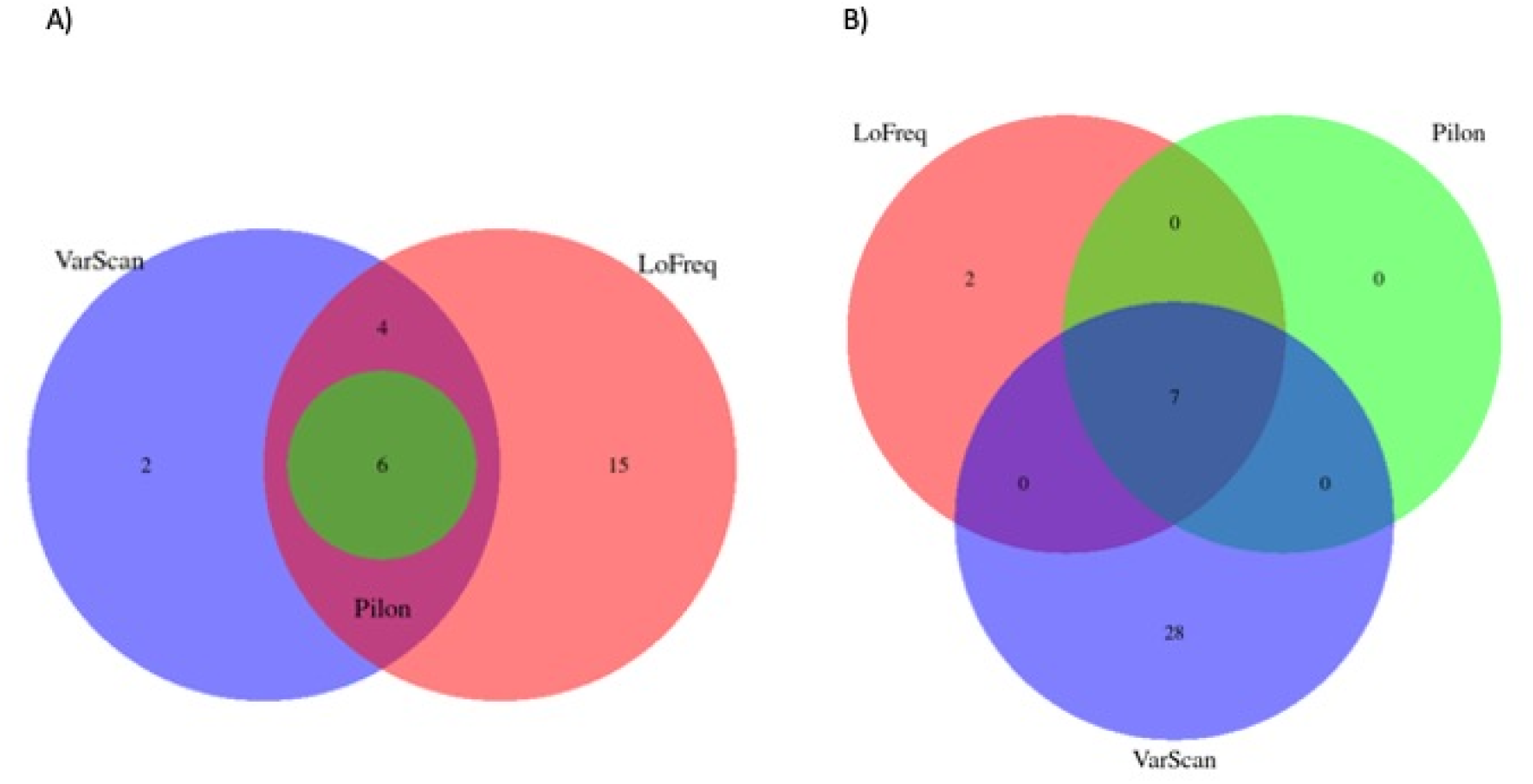
Common and unique SNPs A) for R9 set B) for R10 set

**Fig. 6.**
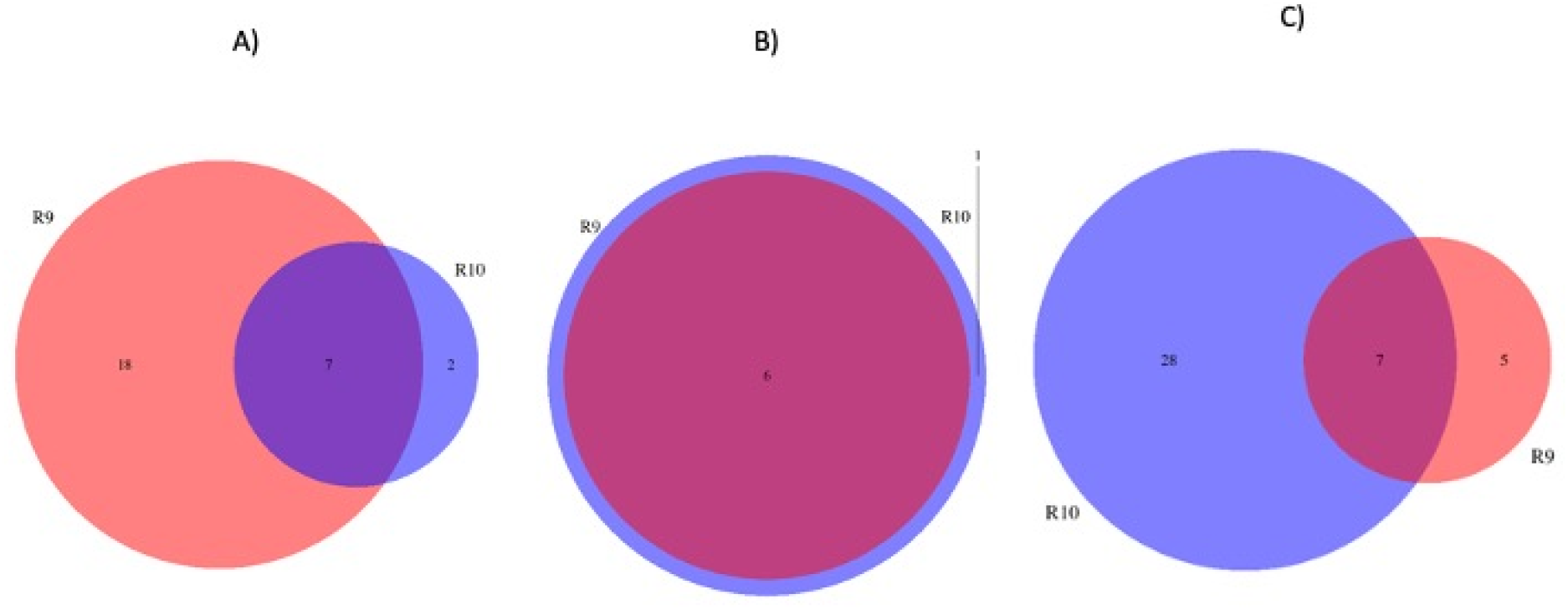
Common and unique SNPs detected by A) LoFreq B) Pilon C) VarScan

**Fig. 7.**
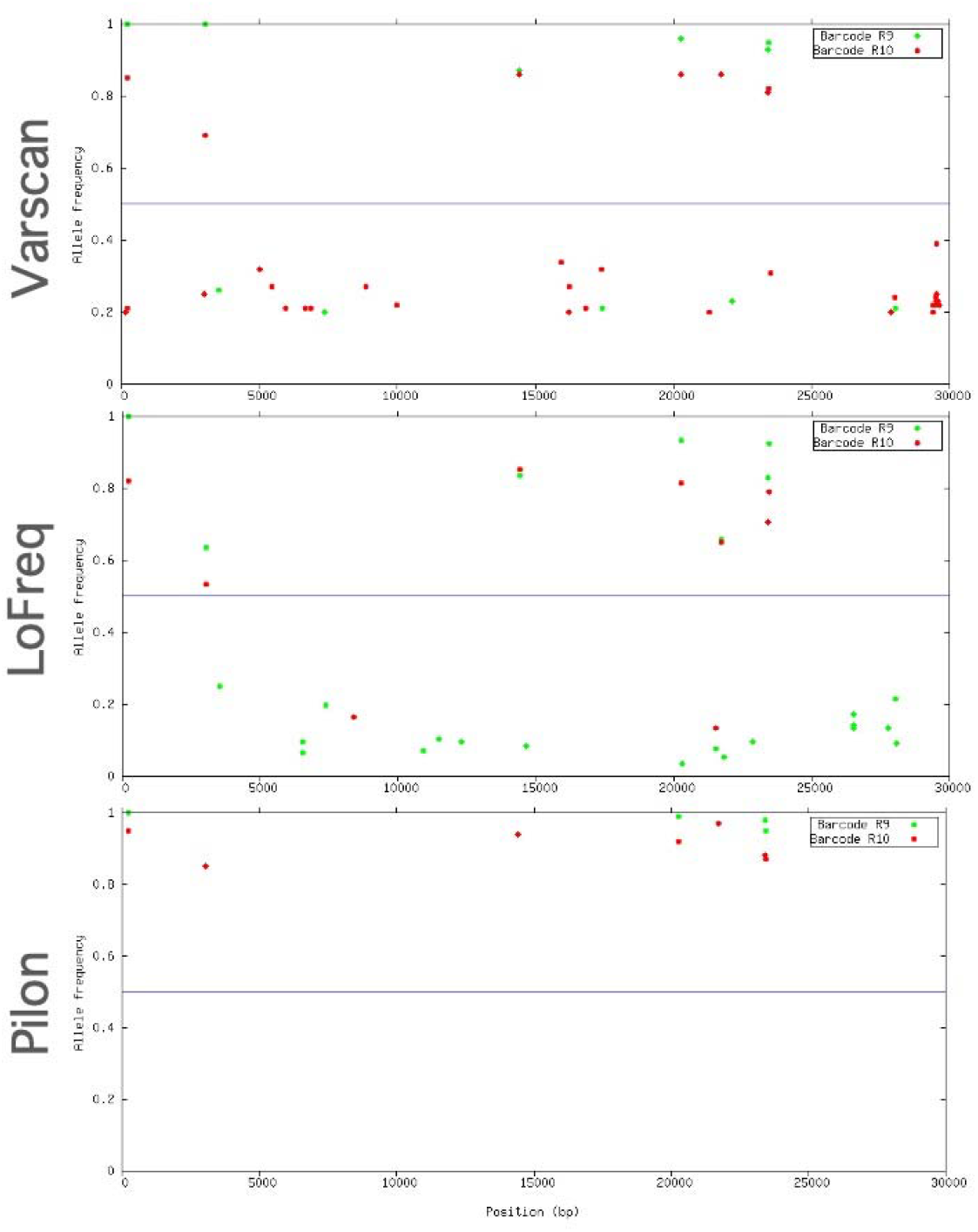
Allele frequencies of SNPs located in the SARS-CoV-2 genome detected by VarScan, LoFreq and Pilon.

**Fig. 8.**
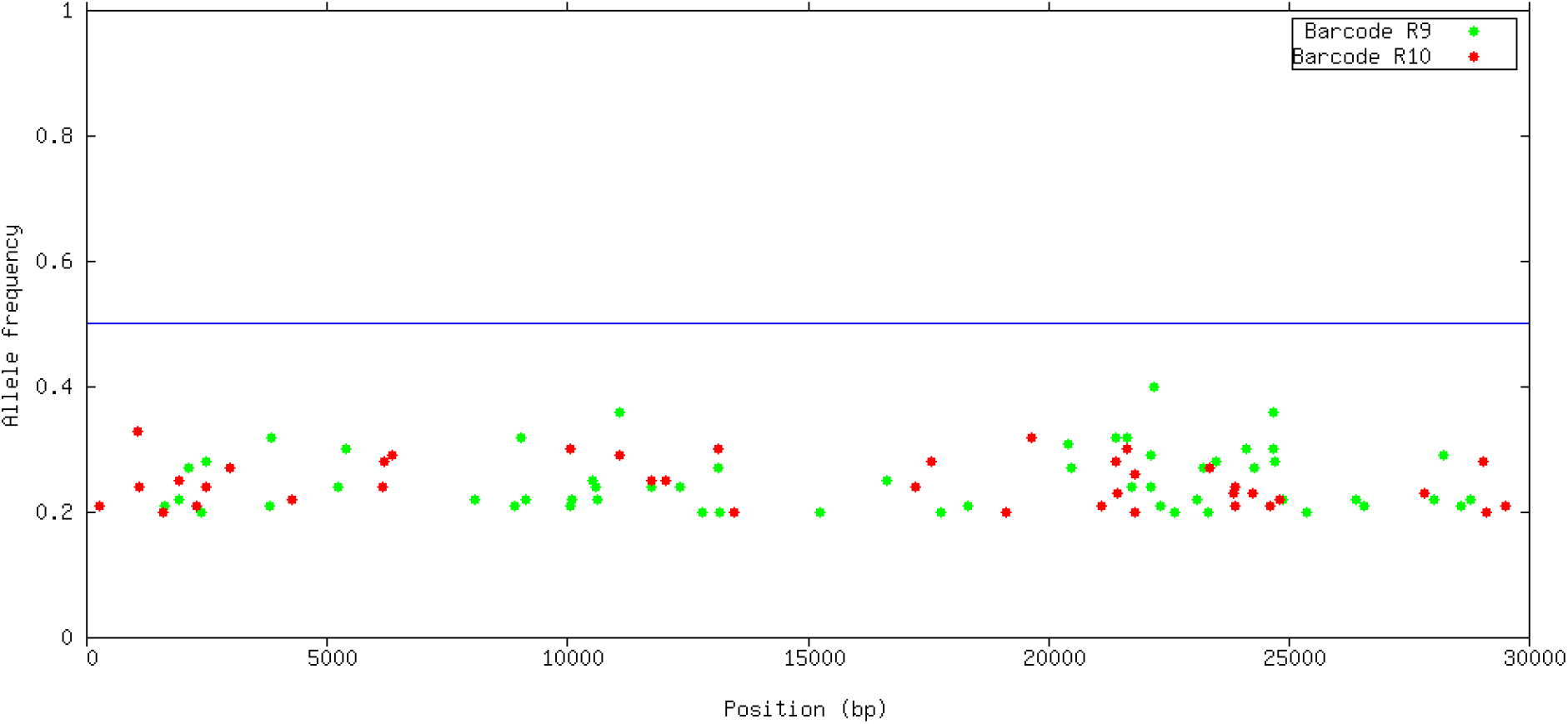
Allele frequencies of INDELs detected by VarScan located in the SARS-CoV-2 genome.

The consensus sequences from both FC were introduced in the pangolin website tool (Rambaut et al. 2020). Both consensus reads belonged to the B1 linage (Boni et al. 2020), which carries the D614G mutation, which is thought to arise in Italy at the beginning of the pandemic in Europe, and spread across Europe and later overseas. This mutation has been related with larger infectivity (Korber et al. 2020).

### De-novo assembly

Fig. 9 and 10 show the dotplot comparison between the SARS-CoV-2 reference genome and the assembly from R9 and R10 FC. Note that R9 assembly is much shorter and sparse than R10 assembly, with a larger number of contigs.

**Fig. 9.**
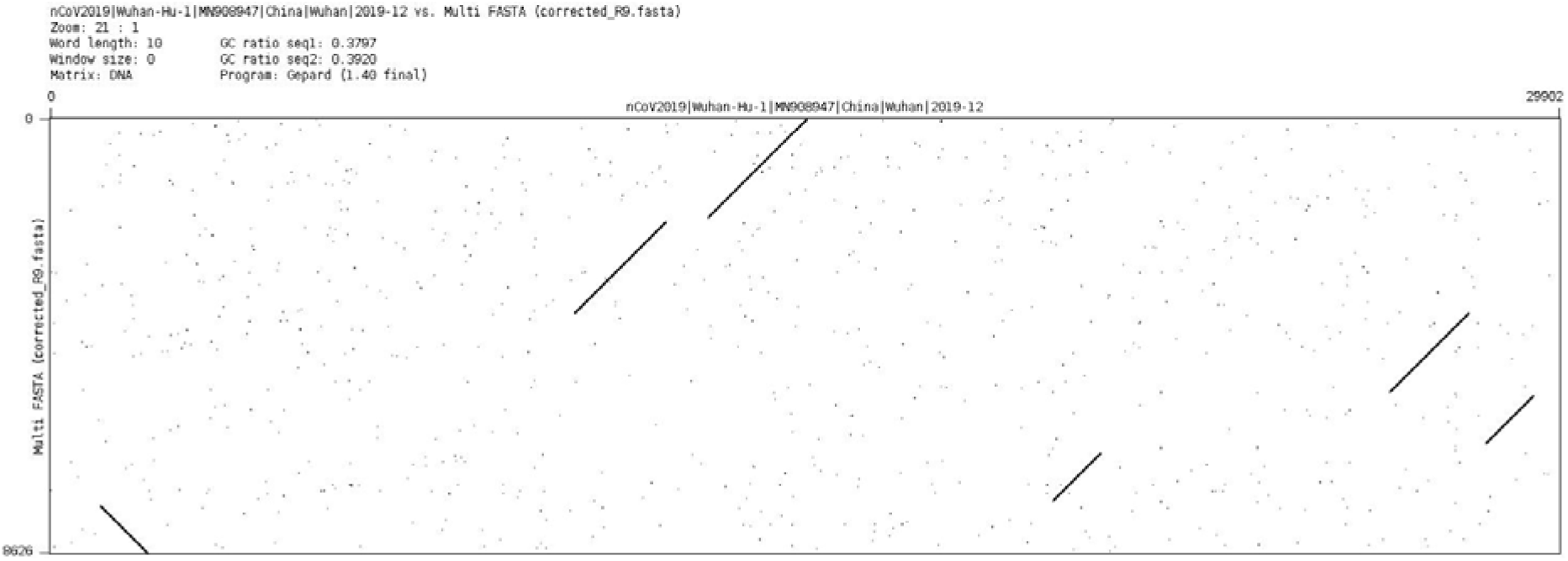
Dotplot comparison of SARS-CoV-2 reference (x-axis) vs. R9 assembly (y-axis)

**Fig. 10.**
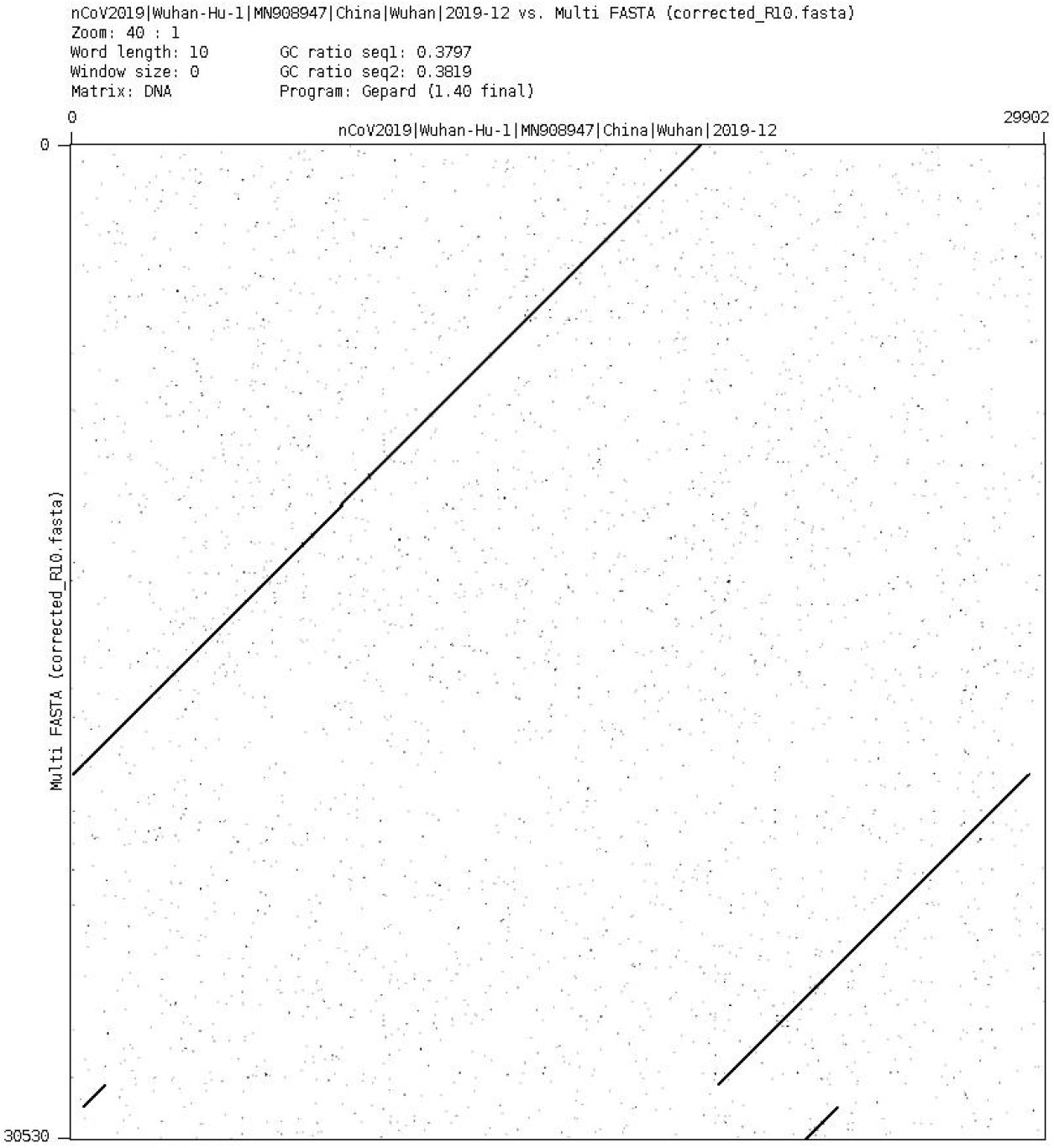
Dotplot comparison of SARS-CoV-2 reference (x-axis) vs. R10 assembly (y-axis)

## DISCUSSION

A clinical sample with RT-qPCR Ct=19.84 for SARS-CoV-2 was used in this study to amplify and sequence the coronavirus genome using Nanopore long reads. Two chemistries and two different protocols were used in the study. A library prepared with the Rapid Barcoding kit (SQK-RBK004) was sequenced on an R9 FC, whereas a library prepared with the Ligation Sequencing Kit (SQK-LSK109) was sequenced on an R10 FC. Both runs yielded similar coverage of the SARS-CoV-2 reference genome. The R9 FC showed a smaller number of mismatches against the reference genome. The Rapid Barcoding kit resulted on shorter sequences as expected, as it uses a transposase fragmentation to insert the barcodes. The alignments showed a large number of variants but most of them in low frequency. Setting a threshold of 0.50 for the variant frequency led to a consistent number of variants per FC and library preparation kit of 6 to 7 variants, regardless the software used. Despite of the larger number of mismatches from R10, the consensus sequence resulted in the same variants detected as from R9.

A de-novo assembly was attempted from both runs. In this case the R10 FC yielded a more complete genome than the R9 run. We interpret that the larger size of the fragments and the larger number of Mb obtained explained this better behaviour, which is mainly due to the library preparation with the Ligation Sequencing Kit rather than to the R10 chemistry.

All called mutations were already described in the GISAID database by the date the sample was collected, except a mutation at position 21727 with a C:T substitution on the S protein, which implies an amino-acid change S:P. This mutation was not observed further in the databases, hence we hypothesis that either this is a sequence error, or the transmission of this mutation was unsuccessful due to the strict lockdown imposed in Madrid from March to May 2020.

It must be pointed out that the effect of the protocol was nested with the FC chemistry. Nonetheless, we were able to extract some conclusions regarding the benefits and drawbacks from each chemistry and library preparation protocols regarding their convenience to sequence the SARS-CoV-2 genome. The R10 chemistry does not improve the quality of the sequenced SARS-CoV-2 genome, and the Ligation Sequencing Kit yielded a much better genome assembly which is recommended for whole genome sequencing of the coronavirus, mainly at low initial DNA concentrations as in the case of this study.

## DECLARATIONS

### Ethics approval

Not applicable

### Consent to participate

Samples were provided under a consent to participate statement.

### Data availability statement

The data that support the findings of this study are available from Seq-Covid consortium (http://seqcovid.csic.es/) who will upload the sequence to the GISAID database. Data are however available from the authors upon reasonable request.

## Authors contribution

P.A.S., C.C.G., J,F.P. extracted the RNA, and performed the cDNA transformation. O.G.R. and M.G.R and C.G. prepared the libraries and performed the cDNA sequencing. R.P.P. developed the computational pipelines for the assemblies and assisted on its analyses. O.G.R., J.F.P, M.J.C. conceived the study and designed the experiments. O.G.R. wrote the manuscript. All authors helped writing and configuring the last version of the manuscript.

## Acknowledgements

We are grateful to Madrid City Council for giving permission to use the samples in this particular study. We greatly thank to all technical and scientific personnel of INIA-CISA who worked voluntarily during the Spanish lockdown conducting mass diagnosis of samples from the essential operational sectors of Madrid City.

## References

Boni MF, Lemey P, Jiang X, Lam TTY, Perry BW, Castoe TA, Rambaut A, Robertson DL (2020) Evolutionary origins of the SARS-CoV-2 sarbecovirus lineage responsible for the COVID-19 pandemic. Nat Microbiol. https://doi.org/10.1038/s41564-020-0771-4

Corman VM, Landt O, Kaiser M, Molenkamp R, Meijer A, Chu DK, Bleicker T, Brünink S, Schneider J, Schmidt ML, Mulders DG, Haagmans BL, van der Veer B, van den Brink S, Wijsman L, Goderski G, Romette J-L, Ellis J, Zambon M, Peiris M, Goossens H, Reusken C, Koopmans MP, Drosten C (2020) Detection of 2019 novel coronavirus (2019-nCoV) by real-time RT-PCR. Eurosurveillance 25:. https://doi.org/10.2807/1560-7917.ES.2020.25.3.2000045

Huson DH, Albrecht B, Bagci C, Bessarab I, Górska A, Jolic D, Williams RBH (2018) MEGAN-LR: New algorithms allow accurate binning and easy interactive exploration of metagenomic long reads and contigs. Biol Direct. https://doi.org/10.1186/s13062-018-0208-7

Koboldt DC, Zhang Q, Larson DE, Shen D, McLellan MD, Lin L, Miller CA, Mardis ER, Ding L, Wilson RK (2012) VarScan 2: Somatic mutation and copy number alteration discovery in cancer by exome sequencing. Genome Res 22:568–576. https://doi.org/10.1101/gr.129684.111

Korber B, Fischer WM, Gnanakaran S, Yoon H, Theiler J, Abfalterer W, Hengartner N, Giorgi EE, Bhattacharya T, Foley B, Hastie KM, Parker MD, Partridge DG, Evans CM, Freeman TM, de Silva TI, Angyal A, Brown RL, Carrilero L, Green LR, Groves DC, Johnson KJ, Keeley AJ, Lindsey BB, Parsons PJ, Raza M, Rowland-Jones S, Smith N, Tucker RM, Wang D, Wyles MD, McDanal C, Perez LG, Tang H, Moon-Walker A, Whelan SP, LaBranche CC, Saphire EO, Montefiori DC (2020) Tracking Changes in SARS-CoV-2 Spike: Evidence that D614G Increases Infectivity of the COVID-19 Virus. Cell 182:812–827.e19. https://doi.org/10.1016/j.cell.2020.06.043

Koren S, Walenz BP, Berlin K, Miller JR, Bergman NH, Phillippy AM (2017) Canu: scalable and accurate long-read assembly via adaptive k-mer weighting and repeat separation. Genome Res 27:722–736. https://doi.org/10.1101/gr.215087.116

Lahti L, Shetty S (2012) microbiome R package

Li H (2016) Minimap and miniasm: fast mapping and de novo assembly for noisy long sequences. Bioinformatics 32:2103–2110. https://doi.org/10.1093/bioinformatics/btw152

McMurdie PJ, Holmes S (2013) phyloseq: An R Package for Reproducible Interactive Analysis and Graphics of Microbiome Census Data. PLoS One 8:e61217. https://doi.org/10.1371/journal.pone.0061217

Oksanen AJ, Blanchet FG, Friendly M, Kindt R, Legendre P, Mcglinn D, Minchin PR, Hara RBO, Simpson GL, Solymos P, Stevens MHH, Szoecs E (2019) Package ‘ vegan ‘

Quick J (2020) nCoV-2019 sequencing protocol. In: https://doi.org/10.17504/protocols.io.bbmuik6w. https://doi.org/10.17504/protocols.io.bbmuik6w. Accessed 20 Sep 2020

Quinlan AR, Hall IM (2010) BEDTools: A flexible suite of utilities for comparing genomic features. Bioinformatics 26:841–842. https://doi.org/10.1093/bioinformatics/btq033

Rambaut A, Holmes EC, O’Toole Á, Hill V, McCrone JT, Ruis C, du Plessis L, Pybus OG (2020) A dynamic nomenclature proposal for SARS-CoV-2 lineages to assist genomic epidemiology. Nat Microbiol 1–14. https://doi.org/10.1038/s41564-020-0770-5

Tillett R, Sevinsky J, Hartley P, Kerwin H, Crawford N, Gorzalski A, Laverdure C, Verma S, Rossetto C, Jackson D, Farrell M, Van Hooser S, Pandori M (2020) Genomic Evidence for a Case of Reinfection with SARS-CoV-2. SSRN Electron J. https://doi.org/10.2139/ssrn.3680955

To KK-W, Hung IF-N, Ip JD, Chu AW-H, Chan W-M, Tam AR, Fong CH, Yuan S, Tsoi H, Ng AC, Lee LL, Wan P, Tso E, To W-K, Tsang D, Chan K, Huang J, Kok K, Cheng VC-C, Yuen K-Y (2020) COVID-19 re-infection by a phylogenetically distinct SARS-coronavirus-2 strain confirmed by whole genome sequencing. Clin Infect Dis 1–25. https://doi.org/10.1093/cid/ciaa1275

Walker BJ, Abeel T, Shea T, Priest M, Abouelliel A, Sakthikumar S, Cuomo CA, Zeng Q, Wortman J, Young SK, Earl AM (2014) Pilon: An integrated tool for comprehensive microbial variant detection and genome assembly improvement. PLoS One 9:. https://doi.org/10.1371/journal.pone.0112963

Wilm A, Aw PPK, Bertrand D, Yeo GHT, Ong SH, Wong CH, Khor CC, Petric R, Hibberd ML, Nagarajan N (2012) LoFreq: A sequence-quality aware, ultra-sensitive variant caller for uncovering cell-population heterogeneity from high-throughput sequencing datasets. Nucleic Acids Res 40:11189–11201. https://doi.org/10.1093/nar/gks918

